# A Modified Tseng Algorithm Approach to Restoring Thoracic Diseases’ Computerized Tomography Images

**DOI:** 10.1101/2023.12.27.573395

**Authors:** Huzaifa Umar, Abubakar Adamu, Ahmad Hijaz, Dilber Uzun Ozsahin

## Abstract

It is well-known that the Tseng algorithm and its modifications have been successfully employed in approximating zeros of the sum of monotone operators. In this study, we restored various thoracic diseases’ computerized tomography (CT) images, which were degraded with a known blur function and additive noise, using a modified Tseng algorithm. The test images used in the study depict calcification of the Aorta, Subcutaneous Emphysema, Tortuous Aorta, Pneumomediastinum, and Pneumoperitoneum. Additionally, we employed well-known image restoration tools to enhance image quality and compared the quality of restored images with the originals. Finally, the study demonstrates the potential to advance monotone inclusion problem-solving, particularly in the field of medical image recovery.

## 1. Introduction

Medical imaging is essential in the diagnosis and treatment of diseases as it provides direct guidance to medical personnel to cure diseases, and over the past decade, advancements in technology have brought about faster, more accurate, and less invasive medical devices [1]. Mathematical models in medical image restoration are a fundamental component of medical imaging, aiming to acquire high-quality images for clinical use while minimizing costs and risks to patients [2]. Intense data-driven models are highly flexible for extracting valuable information from massive data sets yet lack theoretical foundations [3]. Biomedical computing relies on mathematical models. Image data is fundamental in experimental, clinical, biomedical, and behavioural research [4]. Medical imaging problems, such as Magnetic Resonance Imaging (MRI), can be modelled as inverse problems. A modern and practical methodological approach, which has been widely applied, is based on the assumption that most real-life images have a low-dimensional nature. This method is highly effective and has proven to be successful [5]. The analysis of medical datasets through image processing techniques is a critical aspect of modern medical research. The development of algorithms for either partial or fully automatic analysis is essential in this context [6]. Image enhancement stands as a vital and complex method within the realm of image processing technology. Its fundamental goal is to improve the visual quality of an image or present a more polished representation of the picture.

Thoracic diseases (TD) pose significant health challenges, impacting a considerable number of individuals. Chest X-rays, widely utilized as a diagnostic method, play a crucial role in healthcare and are also referred to as computed tomography [7]. TD encompass a range of serious illnesses and health conditions, many of which exhibit a high prevalence. One illustrative example is pneumonia, which annually afflicts millions of individuals globally. In the United States alone, approximately 50,000 people succumb to pneumonia each year [8]. The chest X-ray (CXR) stands out as a widely used and cost-effective diagnostic instrument for identifying chest and thoracic diseases. Deciphering chest X-rays demands substantial expertise and careful visual scrutiny. Despite radiologists undergoing extensive clinical training and professional guidance, errors can still occur due to the intricate nature of diverse lung lesions and the subtle textural differences present in the images [9]. Precise chest X-ray (CXR) interpretations necessitate expert knowledge and medical experience. There have been extensive endeavours to automatically detect thoracic diseases (TD) using CXR data. However, increasing image volumes and subtle texture changes can lead to errors, even by experienced radiologists [10]. Statistical learning methods, such as Support-vector networks [11, 12], Bayesian classifiers [13, 14], and k-nearest neighbour algorithms [15], are not adept at directly handling high-dimensional pixel-level features in medical images. The patterns of diseases found in chest X-rays are numerous, and their occurrence follows a long-tailed (LT) distribution [16]. Therefore, it is essential to have an efficient and reliable mathematical algorithm that can restore the images and easily detect different types of diseases.

The image restoration can be achieved in various forms, such as image denoising [17], deblurring [18], inpainting’ [19,20], dehazing [21], and de-raining [22]. Numerical simulations should be the best option in solving problems related to medical imaging analysis, as factors such as specular noise can affect the interpretation of the image obtained. Therefore, complex mathematical models using various algorithms are needed to study medical images. In this study, various thoracic diseases computerized tomography (CT) images degraded with known blur function and additive noise were restored using a modified Tseng algorithm. Furthermore, we utilized well-known image restoration tools to enhance image quality and compare the quality of the restored images with the original images.

## 2. Methodology and Results

The thoracic disease patients’ CT scan images used were collected from https://www.cancerimagingarchive.net, and they represent various disease conditions [16].

Mathematical model used for image restoration problem are often formulated using equation (1), and illustrated in Figure 1:

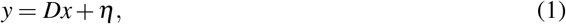

where *y* is the observed image, *D* is the degradation function, *x* is the original image and *η* is noise.

**Figure 1:**
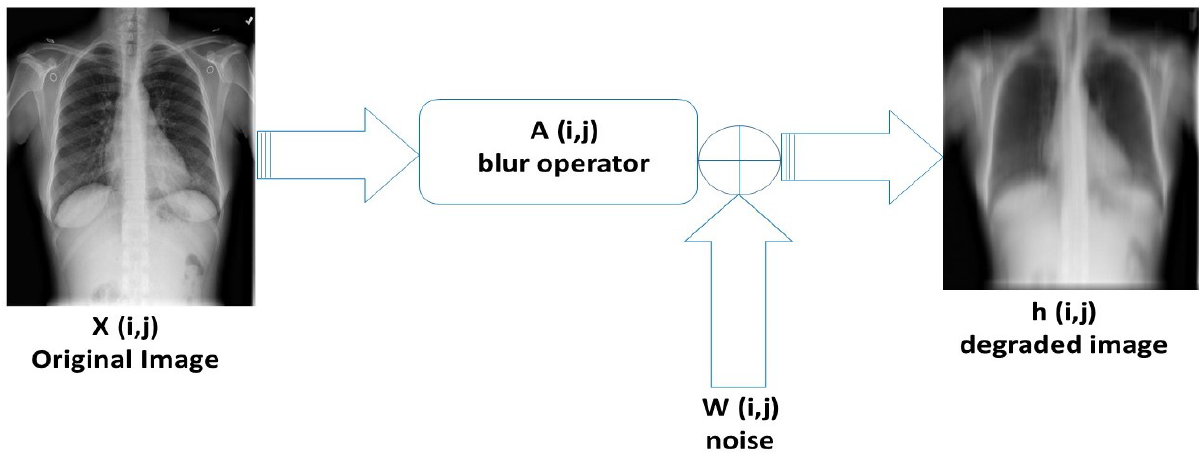
Image Degradation

The objective of this study is to restore a degraded image as illustrated in Figure 1 using mathematical algorithms. Since the solution may not be unique for any degraded image, this problem inherits ill-posedness. To restore well-posedness, regularization techniques are employed. The *l*_1_ regularization method is known to be a powerful technique for image denoising and deblurring problem. The formulation is given by

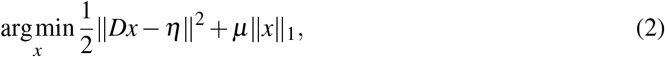

where *μ* is the regularizing term. By simple mathematical reformulation, solutions of the minimization problem (2) are equivalent to solutions of the inclusion problem:

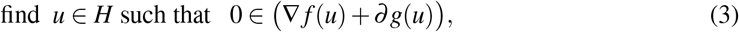

where *H* is a real Hilbert space, ∇ *f* is the gradient of *f* and *∂g* is the subdifferential of *g*, with

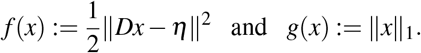

Then,

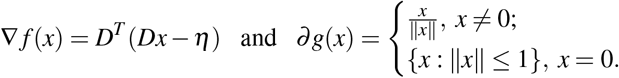

In the literature, several algorithms introduced for approximating zeros of sum of two monotone operators have been used to solve the inclusion problem (3) (see, e.g., [23–26] and the references therein). We propose a modification of the popular Tseng algorithm for approximating solutions of problem (3). Our propose method is the following:

### Algorithm 2.1

**(H) Step 1**. *Given x*_0_ = *Dx* + *η, λ* = 0.001, *μ* = 0.3 *and set k* = 1.

**Step 2**. *Compute w*_*k*_ *and x*_*k*+1_,

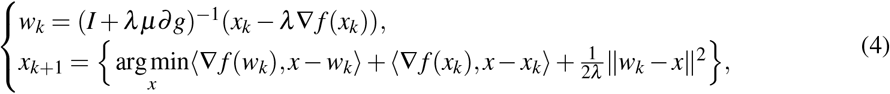

*where I is the identity mapping*.

**Step 3** *Set k ← k* + 1, *and* **go to Step 2**.

### 2.1 Convergence Analysis

#### Theorem 2.2

*The sequence {x*_*k*_*} generated by our proposed method converges to a solution of problem 2*.

**Proof**. Since ∇ *f* is Lipschitz and *∂g* is monotone, the proposed method can be viewed as a corollary of the famous Tseng algorithm [26]. Hence, the convergence analysis follows using a similar argument given in [26]. ▪

### 2.2 Experimental Results and Discussion

In this section, we use the proposed method (Algorithm 2.1) in the restoration process of some CT scan images obtained from thoracic disease patients [16] We label the images as follows: Images 1, 2, 3, 4 and 5 represent Calcification of the Aorta, Subcutaneous Emphysema, Tortuous Aorta, Pneumomediastinum and Pneumoperitoneum, respectively. We study the behaviour and properties of the restored images when they are degraded using MATLAB’s built-in motion blur function (*P* = *f special*(^*′*^*motion*^*′*^, 30, 60)), and we added Gaussian noise (GN) and poison noise (PN) with scaling factor *σ* = 0.001 and 0.05, respectively. The results of the simulations are presented in the Figures 2, 3, 4, 5 and 6 below:

**Figure 2:**
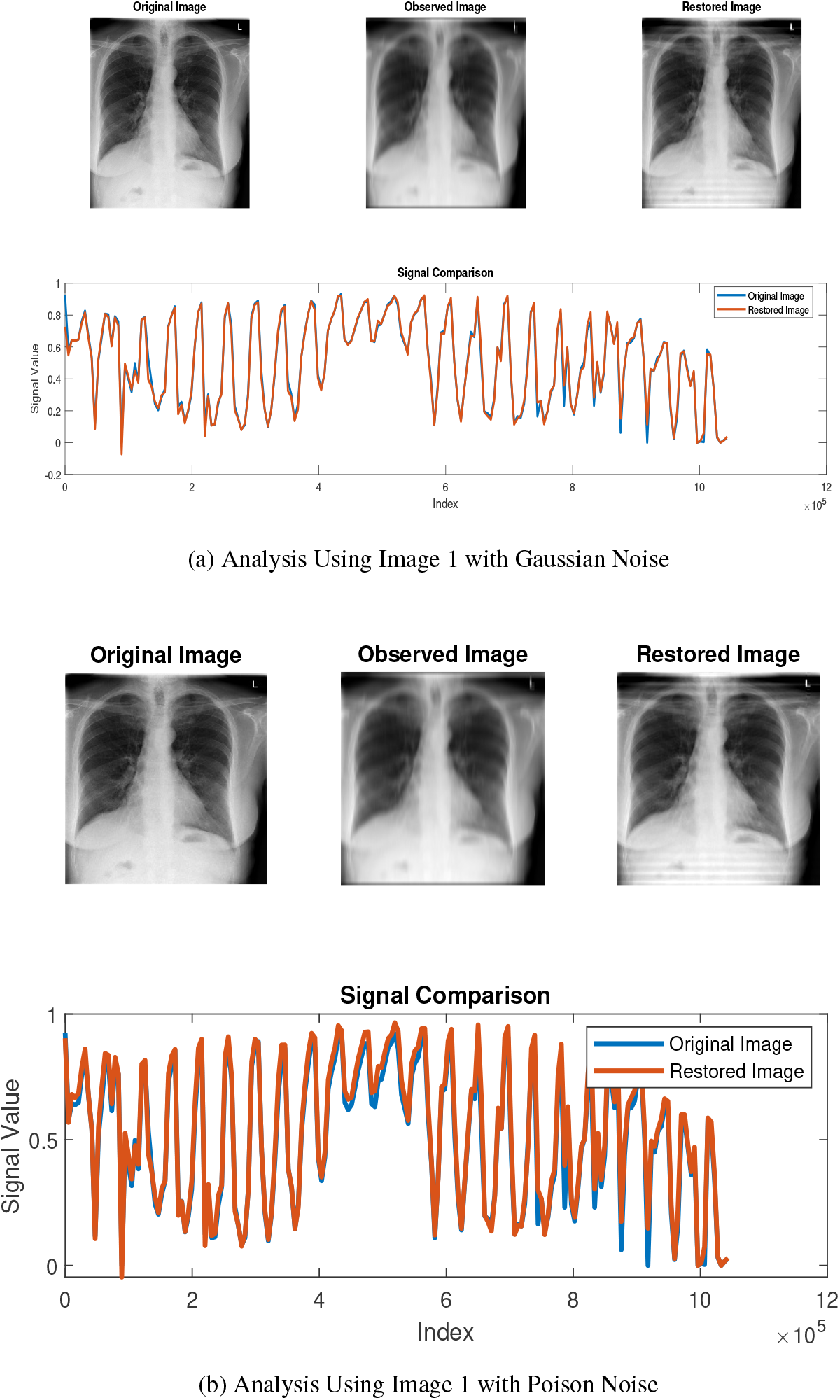
Restoration process via Algorithm 1

**Figure 3:**
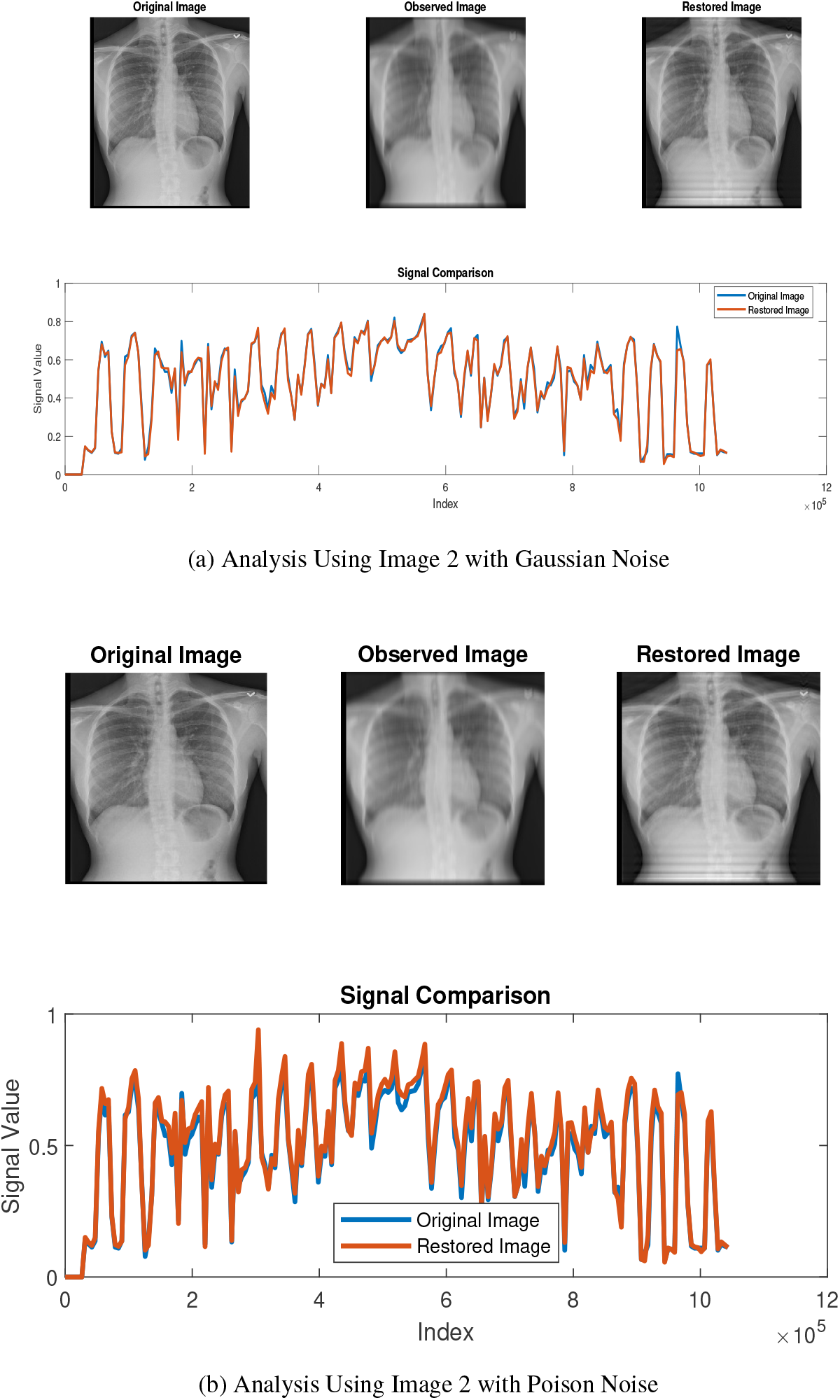
Restoration process via Algorithm 1

**Figure 4:**
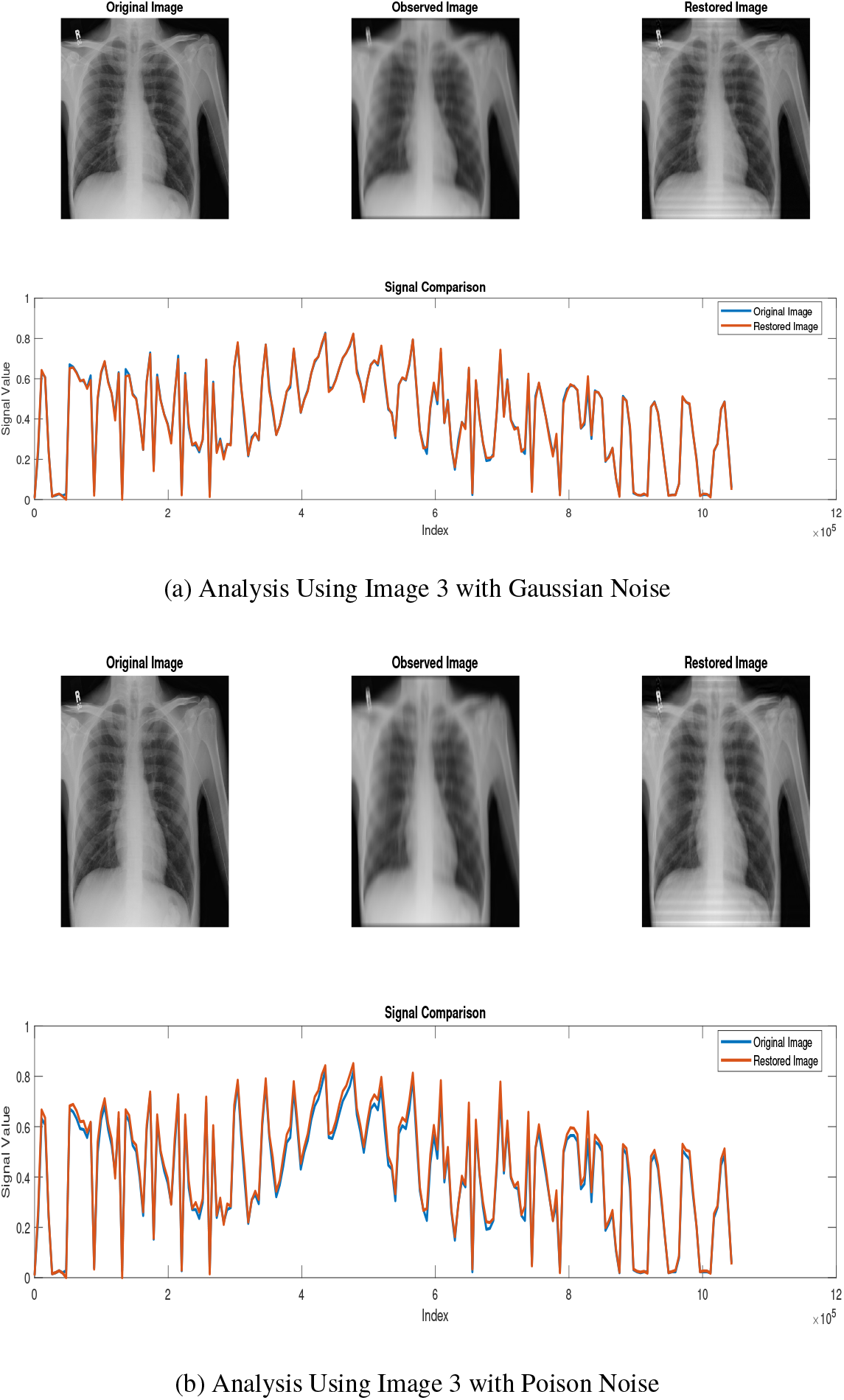
Restoration process via Algorithm 1

**Figure 5:**
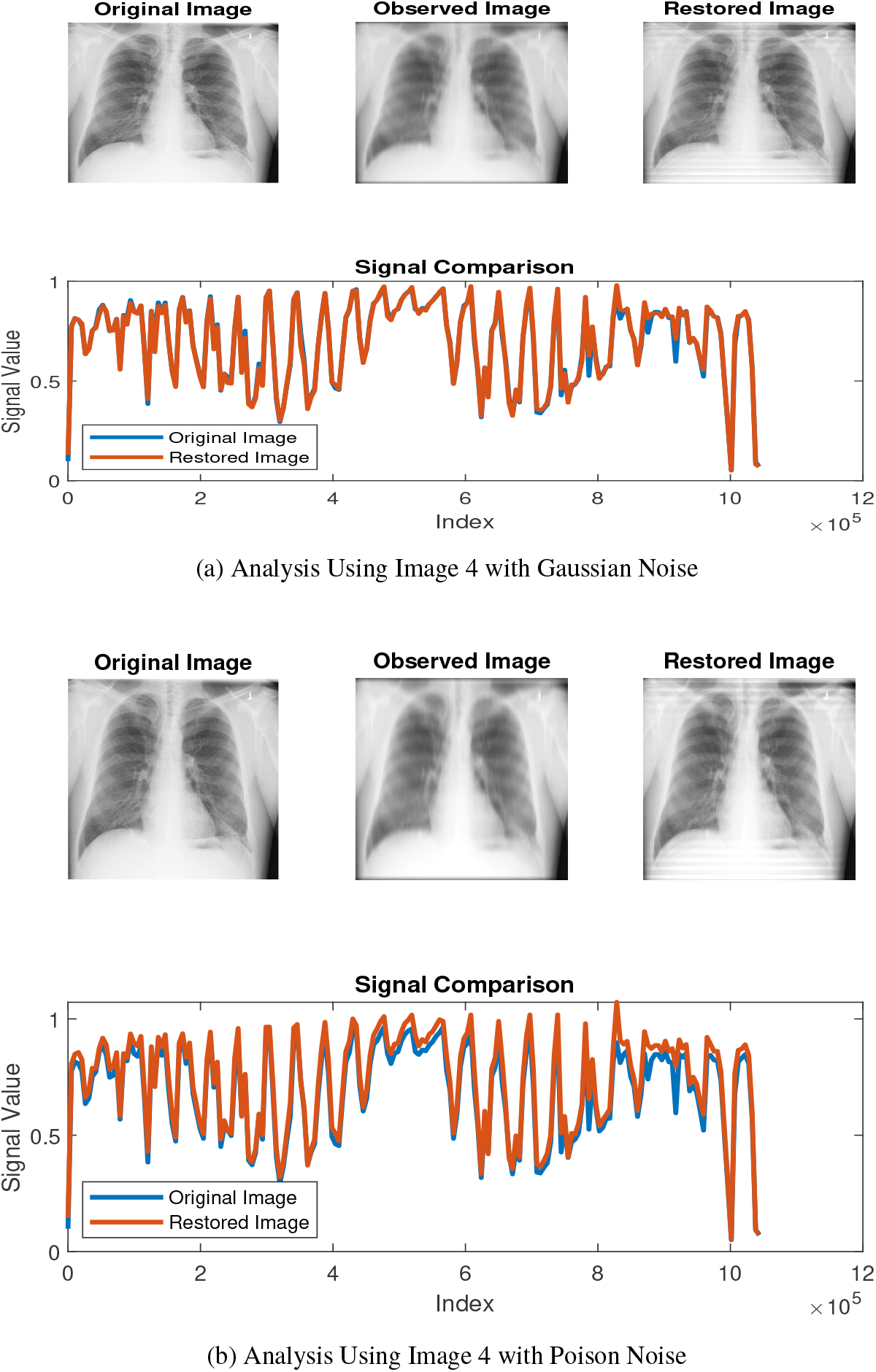
Restoration process via Algorithm 1

**Figure 6:**
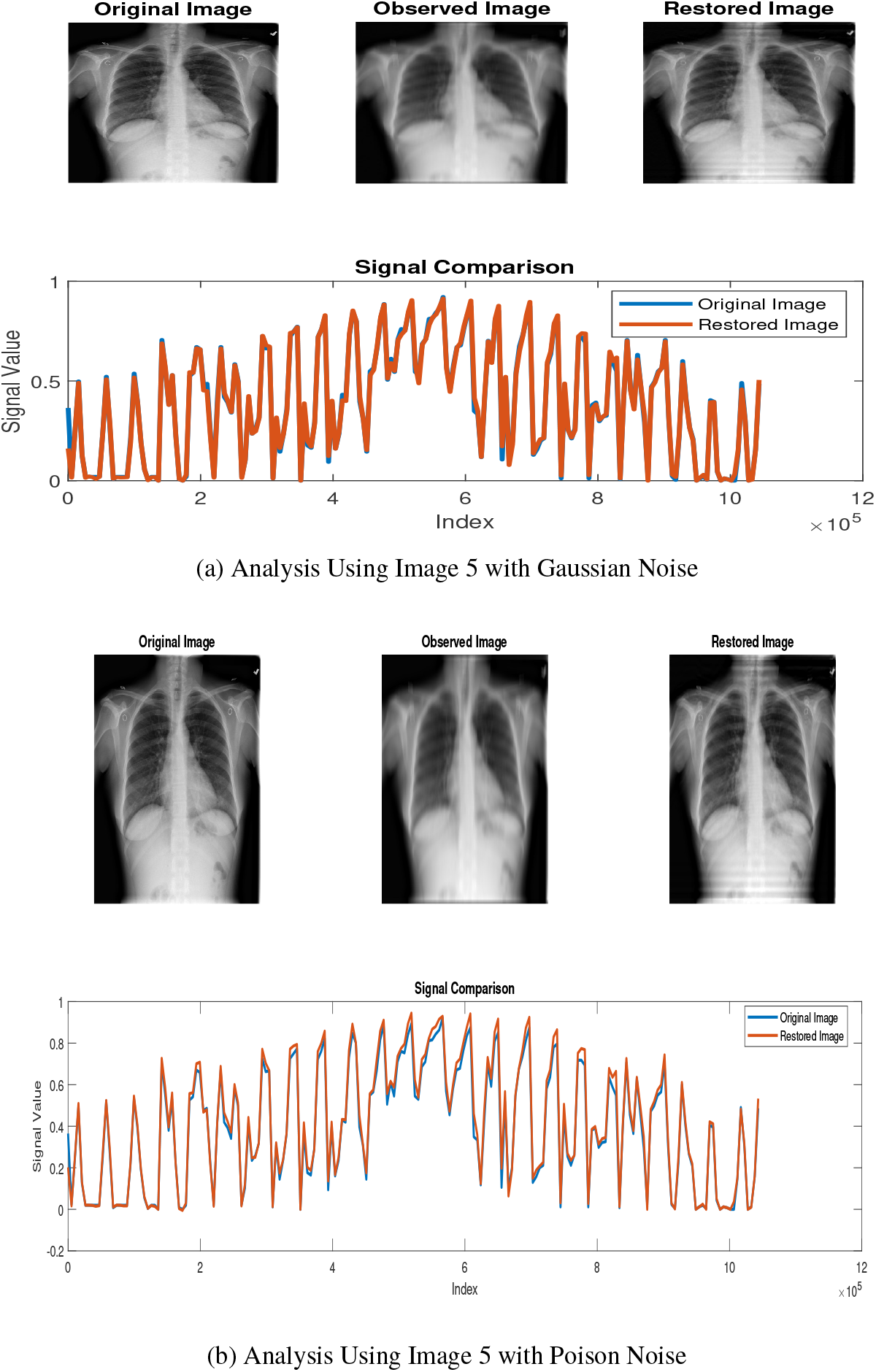
Restoration process via Algorithm 1

## Discussion

Our algorithm was applied to restore computerized tomography (CT) images depicting various thoracic diseases. These images had been intentionally degraded with known blur and additive noise. The implementation of our algorithm led to an enhanced restoration performance for the degraded images. Despite this, the models we introduce exhibit effectiveness specifically in the areas of deblurring and denoising operations, ultimately yielding accurate restoration results.

Clearly, looking at the images in Figures 2, 3, 4, 5 and 6, one can easily see that the proposed Algorithm 1 restored the test images effectively. However, to validate this claim, there are tools use for measuring the quality of restored images. We will use three (3) different metrics to analyze the qualities of the restored images. These tools are namely, the structural dissimilarity index measure (SSIM), the Improvement in signal-to-noise ratio (ISNR) and the signal-to-noise ratio (SNR) index. These metrics are expressed, respectively, as follows:

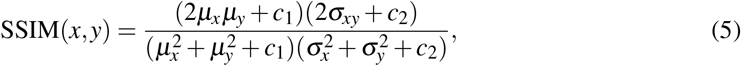

where *x* and *y* represent the original and restored images, *μ*_*x*_ and *μ*_*y*_ denote the mean values of *x* and *y, σ*_*x*_ and *σ*_*y*_ are the standard deviations of *x* and *y, σ*_*xy*_ is the covariance between *x* and *y*, and *c*_1_ and *c*_2_ are small constants introduced to prevent division by zero.

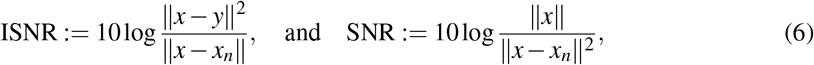

where *x, y*, and *x*_*n*_ denote the original, observed, and estimated images at iteration *n*, respectively.

The SSIM value ranges from 0 to 1, with 1 denoting perfect recovery while higher values for SNR and ISNR indicate superior restoration. The performance of our proposed Algorithm 2.1 using these metrics is detailed in Table 1 below.

**Table 1:**
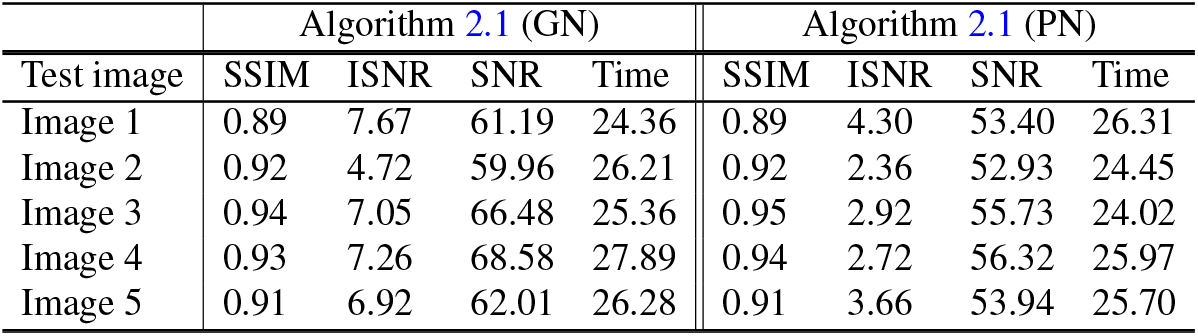
SSIM, ISNR and SNR for the Test Images.

| Test image | Algorithm 2.1 (GN) |  |  |  | Algorithm 2.1 (PN) |  |  |  |
| --- | --- | --- | --- | --- | --- | --- | --- | --- |
|  | SSIM | ISNR | SNR | Time | SSIM | ISNR | SNR | Time |
| Image 1 | 0.89 | 7.67 | 61.19 | 24.36 | 0.89 | 4.30 | 53.40 | 26.31 |
| Image 2 | 0.92 | 4.72 | 59.96 | 26.21 | 0.92 | 2.36 | 52.93 | 24.45 |
| Image 3 | 0.94 | 7.05 | 66.48 | 25.36 | 0.95 | 2.92 | 55.73 | 24.02 |
| Image 4 | 0.93 | 7.26 | 68.58 | 27.89 | 0.94 | 2.72 | 56.32 | 25.97 |
| Image 5 | 0.91 | 6.92 | 62.01 | 26.28 | 0.91 | 3.66 | 53.94 | 25.70 |

### Remark 1

*From the metrics of the restored images presented in Table 1, it is convincing that our proposed Algorithm 2.1 restored the test images with high quality*.

## 3. Conclusion

Overall, this study successfully restored computerized tomography (CT) images of various thoracic diseases that had been degraded with known blur and additive noise. The restoration was achieved through the implementation of a mathematical algorithm, specifically a modified Tseng algorithm. Furthermore, we employed established image restoration tools to enhance the quality of the images and conducted a comprehensive comparison between the restored images and the original ones. This approach not only showcases the efficacy of the applied algorithm but also underscores the importance of combining mathematical models with established tools in the field of medical image restoration.

## 4. Declaration

## 4.1 Acknowledgment

The authors appreciate the support provided by their institutions. “Operational Research Centre for Healthcare, Near East University, TRNC.”

## 4.3 Use of AI

The authors declare that they did not use AI to generate any part of the paper.

## 4.4 Data Availability

All data is included in the manuscript.

## 4.6 Ethical Approval

Not Applicable.

